# A murine experimental model of the unique pulmonary thrombotic effect induced by the venom of the snake *Bothrops lanceolatus*

**DOI:** 10.1101/2024.07.03.601850

**Authors:** Alexandra Rucavado, Erika Camacho, Teresa Escalante, Bruno Lomonte, Julián Fernández, Daniela Solano, Isabel Quirós-Gutiérrez, Gabriel Ramírez-Vargas, Karol Vargas, Ivette Argüello, Alejandro Navarro, Carlos Abarca, Álvaro Segura, Jonathan Florentin, Hatem Kallel, Dabor Resiere, Remi Neviere, José María Gutiérrez

**Affiliations:** Instituto Clodomiro Picado, Facultad de Microbiología, Universidad de Costa Rica, San José, Costa Rica; Laboratorio de Hematología, Hospital Nacional de Niños ‘Dr Carlos Sáenz Herrera’, Caja Costarricense del Seguro Social, San José, Costa Rica; Department of Toxicology and Critical Care Medicine, University Hospital of Martinique (CHU Martinique), Fort-de-France, France; Intensive Care Unit, Cayenne General Hospital, French Guiana; Tropical Biome and immunopathology CNRS UMR-9017, Inserm U 1019, Université de Guyane, French Guiana; Cardiovascular Research Team (UR5_3 PC2E), University of the French West Indies (Université des Antilles), 97200 Fort de France, France

**Keywords:** *Bothrops lanceolatus*, Martinique, thrombosis, coagulopathy, proteome, venom from juvenile specimens

## Abstract

**Background:** The venom of *Bothrops lanceolatus*, a viperid species endemic to the Lesser Antillean Island of Martinique, induces a unique clinical manifestation, i.e., thrombosis. Previous clinical observations indicate that thromboses are more common in patients bitten by juvenile specimens. There is a need to develop an experimental model of this effect in order to study the mechanisms involved.

**Methodology/principal findings:** The venoms of juvenile and adult specimens of *B. lanceolatus* were compared by (a) describing their proteome, (b) assessing their ability to induced thrombosis in a mouse model, and (c) evaluating their *in vitro* procoagulant activity and *in vivo* hemostasis alterations. Venom proteomes of juvenile and adult specimens were highly similar. When injected by the intraperitoneal (i.p.) route, the venom of juvenile specimens induced the formation of abundant thrombi in the pulmonary vasculature, whereas this effect was less frequent in the case of adult venom. Thrombosis was not abrogated by the metalloproteinase inhibitor Batimastat. Both venoms showed a weak *in vitro* procoagulant effect on citrated mouse plasma and bovine fibrinogen. When administered intravenously (i.v.) venoms did not affect classical clotting tests (prothrombin time and activated partial thromboplastin time) but caused a partial drop in fibrinogen concentration. The venom of juvenile specimens induced partial alterations in some rotational thromboelastometry parameters after i.v. injection. No alterations in coagulation tests were observed when venoms were administered i.p., but juvenile and adult venoms induced a marked thrombocytopenia.

**Conclusions/significance:** An experimental model of the thrombotic effect induced by *B. lanceolatus* venom was developed. This effect is more pronounced in the case of venom of juvenile specimens, despite the observation that juvenile and adult venom proteomes are similar. Adult and juvenile venoms do not induce a consumption coagulopathy characteristic of other *Bothrops* sp venoms. Both venoms induce a conspicuous thrombocytopenia. This experimental model reproduces the main clinical findings described in these envenomings and should be useful to understand the mechanisms of this thrombotic effect.

**Author summary:** Envenomings by the viperid species *Bothrops lanceolatus*, endemic of the Caribbean Island of Martinique, are characterized by a unique thrombotic effect responsible for infarcts in various organs. Until now, no experimental *in vivo* models of this effect have been described. In this study, we developed a mouse model of thrombosis by using the intraperitoneal route of venom injection. The venom of juvenile specimens of *B. lanceolatus* induced the formation of abundant thrombi in the lungs, whereas the effect was much less pronounced with the venom of adult specimens. This difference in the ability of juvenile and adult venoms occurs despite both venoms having highly similar proteomic profiles. Both adult and juvenile venoms showed a weak *in vitro* procoagulant effect on plasma and fibrinogen, underscoring a thrombin-like (pseudo-procoagulant) activity. *In vivo*, the venoms did not affect the classical clotting tests (prothrombin time and activated partial thromboplastin time) but induced a partial drop in fibrinogen concentration and limited alterations in rotational thromboelastometry parameters when injected by the i.v. route. In contrast, few alterations of these parameters were observed after i.p. injection of venoms, in conditions in which thrombosis occurred, hence evidencing the lack of a consumption coagulopathy. After i.p. injection both venoms induced a pronounced thrombocytopenia. This experimental model reproduces some of the main clinical manifestations of envenoming by this species. This model can be used to identify the toxins responsible for the thrombotic effect, to study the mechanism(s) of thrombosis and to assess the preclinical efficacy of antivenoms.

## Introduction

*Bothrops lanceolatus* is a viperid snake species endemic to the Lesser Caribbean Island of Martinique as a result of a long-distance dispersal of South American species of the *Bothrops atrox-asper* complex [1]. It inflicts between 20-30 cases of envenoming per year [2, 3]. The clinical manifestations of these envenomings include local effects, i.e., pain, edema, and necrosis, and systemic alterations, i.e., hemodynamic disturbances and, in some cases, alterations in hemostasis [2–5], most of which are characteristic of envenomings by *Bothrops* sp [6–8]. However, in contrast to the typical hemostatic alterations induced by *Bothrops* sp venoms, characterized by a consumption coagulopathy and defibrinogenation [8, 9], envenomings by *B. lanceolatus*, in the absence of antivenom treatment, involve a severe thrombotic effect which may result in cerebral, myocardial, and pulmonary infarctions [3, 10, 11], and a diffuse thrombotic microangiopathy [12]. Interestingly, clotting laboratory tests are altered to a much lower extent in these envenomings as compared to those inflicted by other *Bothrops* species, although thrombocytopenia is frequent [3, 5, 11]. This thrombotic effect has been also described in envenomings by the closely relates species *B. caribbaeus*, endemic to the neighboring island of Saint Lucia [13].

Despite its clinical relevance, the pathogenic mechanisms and the toxins involved in this unique thrombotic effect remain unknown. It has been proposed that venom-induced alterations in the endothelium might be involved [12, 13, 14]. Other proposed mechanisms include platelet activation, the effect of venom on the binding of von Willebrand factor to type VI collagen in the subendothelium [15], and the proinflammatory activity of the venom [16–18].

Proteomic analysis of adult specimens of *B. lanceolatus* venom have revealed a pattern characteristic of viperid snake venoms, with predominance of P-III and P-I metalloproteinases (SVMPs), serine proteinases (SVSPs), phospholipases A_2_ (PLA_2_s) and, to a lower extent, L-amino acid oxidases and C-type lectin like proteins (SNACLECs), disintegrins, and cysteine-rich secretory proteins (CRISPs) [14, 19]. The proteome and the toxicological profile of the venom of juvenile specimens have not been investigated. Experimental *in vitro* and *in vivo* studies with venom of adult specimens have documented lethal, hemorrhagic, edema-forming, myotoxic, PLA_2_, proteinase, fibrinogenolytic, complement-activating, and proinflammatory activities [14, 16–18, 20, 21, 22]. Conflicting results have been presented regarding the *in vitro* procoagulant activity of this venom on plasma, since this effect has been observed in some studies [19, 23, 24] but not in others [20, 21]. On the other hand, the thrombotic effect described in humans has not been reproduced in mice injected intravenously (i.v.) with venom [14]. Thus, there is a need to develop experimental models which would allow the study of the mechanisms involved in this thrombotic effect.

It has been described that the thrombotic effect is more frequently observed in patients bitten by small, i.e., juvenile snakes [4, 25], thus raising the possibility of ontogenetic variations in the composition and effects of the venom of this species. In order to develop an experimental model of venom-induced thrombosis, venoms were collected from juvenile and adult specimens to compare their proteomes and their effects on blood coagulation, platelet numbers, and thrombi formation. Results revealed that the venom of juvenile specimens, when injected intraperitoneally, induces thrombosis in the pulmonary vasculature, whereas the venom of adult specimens has a much weaker thrombotic activity. This thrombotic effect occurs in the absence of a consumption coagulopathy characteristic of other *Bothrops* sp venoms.

## Methods

### Ethical statement

The methods carried out in this study using mice were approved by the Institutional Committee for the Care and Use of Laboratory Animals of Universidad de Costa Rica (approval code CICUA 13-2023). Mice (CD-1 strain, 20-23 g body weight of both sexes) were provided by the Bioterium of Instituto Clodomiro Picado. Mice were handled in Tecniplast Eurostandard Type II 1264C cages (268 x 215 x 141 mm), four mice per cage. Mice were maintained at 18 – 24°C, 60 – 65% relative humidity, and a 12:12 h light-dark cycle, and were provided with water and food *ad libitum*.

### Venoms

Venoms were obtained from *Bothrops lanceolatus* specimens caught in the wild in various locations of Martinique. Venom of adult snakes was a pool obtained from six specimens (five females, one male) with a range of body length of 150 to 172 cm. Venom of juvenile snakes was a pool prepared from ten specimens (five females, five males) with a range of body length of 59 to 92 cm. After venom extraction, snakes were released to the field in the places where they had been collected. Upon collection, venoms were frozen, freeze dried, and kept at -40 °C until used. Solutions of venoms were prepared immediately before use.

### SDS-Polyacrylamide gel electrophoresis (SDS-PAGE)

The venoms of *B. lanceolatus* (adults and juveniles) were comparatively analyzed by SDS-PAGE, after reduction with 2-mercaptoethanol for 5 min at 95°C. Samples of 10, 20 or 40 µg were loaded onto a 4-20% pre-cast gel (Bio-Rad) alongside with molecular weight markers (Bio-Rad) and separated at 160 v in a mini-Protean apparatus (Bio-Rad). Proteins were visualized with Coomassie Blue R-250 stain and recorded with ImageLab® software

### Reverse phase HPLC (RP-HPLC)

The venoms of *B. lanceolatus* (adults and juveniles) were comparatively analyzed by RP-HPLC. Samples of 2 mg were dissolved in 200 µL of water containing 0.1% trifluoroacetic acid (solution A) and centrifuged. The supernatant was applied to a reverse-phase column (C_18_, 250 x 4.6 mm, 5 µm particle; Phenomenex) equilibrated with the same solution and separated at 1 mL/min using an Agilent 1220 chromatography system monitored at 215 nm. Elution was carried out with a gradient toward acetonitrile containing 0.1% trifluoroacetic acid, as follows: 0% for 5 min, 0-15% for 10 min, 15-45% for 60 min, 45-70% for 10 min, and 70% for 9 min [26].

### Proteomic profiling

The venoms of *B. lanceolatus* (adults and juveniles) were comparatively analyzed using a bottom-up ’shotgun’ MS/MS approach, as described [27]. In brief, 15 μg of each venom, dissolved in 25 mM ammonium bicarbonate, were subjected to reduction with 10 mM dithiothreitol (30 min at 56 °C), alkylation with 50 mM iodoacetamide (20 min in the dark), and overnight digestion with sequencing grade trypsin at 37 °C. After stopping the reaction with 0.5 μL of formic acid, the tryptic peptides were dried in a vacuum centrifuge (Eppendorf), redissolved in 0.1% formic acid, and separated by RP-HPLC on a nano-Easy 1200^®^ chromatograph coupled to a Q-Exactive Plus^®^ mass spectrometer (Thermo). Six μL of each sample (∼0.7 μg of peptide mixture) were loaded onto a C18 trap column (75 μm × 2 cm, 3 μm particle; Thermo), washed with 0.1% formic acid (solution A), and separated at 200 nL/min on a C18 Easy-Spray PepMap^®^ column (75 μm × 15 cm, 3 μm particle; Thermo). A gradient toward solution B (80% acetonitrile, 0.1% formic acid) was developed for a total of 120 min (1–5% B in 1 min, 5–26% B in 84 min, 26–80% B in 30 min, 80–99% B in 1 min, and 99% B for 4 min). MS spectra were acquired in positive mode at 1.9 kV, with a capillary temperature of 200 °C, using 1 μscan in the range 400–1600 m/z, maximum injection time of 50 msec, AGC target of 1×10^6^, and resolution of 70,000. The top 10 ions with 2–5 positive charges were fragmented with AGC target of 3×10^6^, minimum AGC 2×10^3^, maximum injection time 110 msec, dynamic exclusion time 5 s, and resolution 17,500. MS/MS spectra were processed against protein sequences contained in the UniProt database for Serpentes (taxid:8570) using Peaks X^®^ (Bioinformatics Solutions). Parent and fragment mass error tolerances were set at 15.0 ppm and 0.5 Da, respectively. Cysteine carbamidomethylation was set as fixed modification, while methionine oxidation and deamidation of asparagine or glutamine were set as variable modifications. A maximum of 2 missed cleavages by trypsin in semispecific mode were allowed. Filtration parameters for match acceptance were set to FDR<0.1%, detection of ≥1 unique peptide, and -10lgP protein score ≥30.

### *In vitro* coagulant activity on plasma and fibrinogen

Blood was collected from mice by cardiac puncture under isoflurane anesthesia, and immediately added to citrate anticoagulant (3.8% sodium citrate; citrate: blood volume ratio of 1:9), followed by centrifugation at 2,000 x g for 10 min for the collection of plasma. Aliquots of 200 µL of citrated plasma were incubated at 37°C for 5 min, and then 15 µL of 0.2 M CaCl_2_ was added, followed by 25 µL of various amounts of each venom, dissolved in 25 mM Tris-HCl, 137 mM NaCl, 3.4 mM KCl, pH 7.4 (TBS). Tubes were incubated at 37 °C and the clotting time recorded. The ability of venoms to clot fibrinogen was assessed by incubating 200 µL of a 6 mg/mL bovine fibrinogen solution (Merck, Darmstadt, Germany) at 37 °C for 5 min, followed by the addition of 25 µL of TBS containing various amounts of venoms. In both cases, the formation of a clot was visually assessed during a period of 10 min by tilting the tubes every minute. Procoagulant activity was also assessed by the turbidimetric assay described by O’Leary and Isbister [28], as modified by Sánchez et al. [29]. Briefly, 50 µg venom, dissolved in 100 µL TBS, were added to wells in a microplate and incubated for 2 min at 37°C in a microplate reader (Cytation 3 Imaging Reader, BioTek, VT, USA). Then, 4 µL of 0.4 M CaCl_2_ were added to 100 µL of mouse citrated plasma previously incubated at 37°C, and the mixture added to wells in the plate containing the venom dilutions. Controls included plasma/CaCl_2_ incubated with TBS with no venom. After shaking for 5 sec, the absorbances at 340 nm were recorded during 10 min. In other experiments, the same protocol was followed, but a 6 mg/mL solution of bovine fibrinogen (Merck) was used instead of plasma, without the addition of CaCl_2_, in order to assess for thrombin-like (pseudo-procoagulant) activity. In this assay, various amounts of venom were tested and absorbances recorded at 10 min, and a dose of 25 µg venom was tested at various time intervals. In all cases, tests were done in triplicates.

### Effects of venom on coagulation parameters *in vivo*

Groups of four mice (20-22 g) received an intravenous (i.v.) injection of 20 µg of either juvenile or adult *B. lanceolatus* venom, dissolved in 100 µL 0.12 M NaCl, 0.04 M phosphate, pH 7.2 solution (PBS). One hr after injection, mice were bled by cardiac puncture under isoflurane anesthesia and immediately added to Eppendorf vials containing 3.8 % sodium citrate as anticoagulant, using a citrate : blood volume ratio of 1 : 9. In another set of experiments, a dose of 70 µg of either adult or juvenile *B. lanceolatus* venom, dissolved in 100 µL PBS, was injected by the intraperitoneal (i.p.) route into groups of four mice (20-22 g). Blood was collected 4 hr after injection and added to citrate anticoagulant, as described above. This dose was selected because it induced pulmonary thrombosis at this time interval (see below). For controls, groups of mice received an injection of 100 µL PBS by the i.v. route and were bled one hr after injection, as described.

Citrated blood samples were used for rotational thromboelastometry determinations, using a ROTEM Delta 4000 equipment according to the manufacturer’s instructions (Tem Innovations, GmbH, Munich, Germany). The parameters determined were: Extem, Intem and Fibtem clotting time (CT), clot formation time (CFT), and amplitude-clot strength at 20 min (A20). Extem and Intem tests evaluate the extrinsic and intrinsic coagulation pathways, respectively, whereas Fibtem evaluates the contribution of fibrinogen to clot formation and strength in conditions in which platelets are inhibited. CT is the time lapse (in sec) needed for the formation of a clot amplitude of 2 mm. CFT is the time lapse (in sec) between 2 mm clot amplitude and 20 mm clot amplitude. A20 reflects the clot firmness (in mm amplitude) 20 min after CT. These tests were run at a temperature of 37°C. Additional individual citrated blood samples from each experimental group were centrifuged at 2,000 g for 10 min, and plasma was obtained for determination of prothrombin time (PT), activated partial thromboplastin time (aPTT) and fibrinogen concentration, using a STA R Max2 coagulation analyzer (Stago, Paris, France). For platelet counts, a dose of 70 µg of either adult or juvenile *B. lanceolatus* venom, dissolved in 100 µL PBS, was injected i.p. into groups of four to eleven mice (20-22 g), whereas a control group of seven mice received 100 µL PBS alone. Blood was collected at 4 hr and added to sodium citrate as described. Platelet counts were carried out in citrated blood in an automated hematology analyzer (VetsCan HM5, Abaxis Global diagnostics, USA). The time of 4 hr was selected in the case of mice receiving an i.p. injection of venom because thrombi developed in the lungs at this time interval (see below).

### Histological assessment of thrombotic activity

In order to establish a model of the thrombotic activity described in patients envenomed by *B. lanceolatus*, various routes of venom injection were initially tested. Groups of four mice (22-23 g) received either 30 µg venom by the i.v. route in the caudal vein, 50 µg venom by the i.m. route in the right gastrocnemius muscle, 50 µg by the subcutaneous (s.c.) route or 70 µg by the i.p. route, in all cases diluting the venom in 100 µL PBS. Groups of four control mice received 100 µL PBS under otherwise identical conditions. These venom doses were selected for being sublethal (in the case of i.v. and i.p. routes) on the basis of previous reports of Median Lethal Dose (LD_50_) of venom from adult specimens [21, 30], and for inducing prominent local tissue pathology without being lethal (in the cases of i.m. and s.c. routes). At either 4 or 24 hr after venom injection mice were sacrificed by cervical dislocation, and samples of heart, brain and lungs were obtained and added to 3.7% formalin fixative solution.

Tissues were processed routinely and embedded in paraffin. Sections of 4 µm were obtained and stained with hematoxylin-eosin for microscopic observation.

### Staining for fibrin in the microvasculature

In order to ascertain whether fibrin microthrombi developed in the pulmonary microvasculature, paraffin-embedded sections from mice receiving i.p. injections of 70 µg of venom of juvenile specimens were prepared as described above and stained with the Martius-Scarlet-Blue kit for staining fibrin (Diapath, Martinengo, Italy). Sections of 4 µm were deparaffinized with xylene and ethanol and rehydrated with water. Then, sections were serially stained with martius yellow, crystal scarlet and methyl blue following the manufacturer’s instructions. Sections were briefly rinsed with 1% acetic acid, dehydrated and mounted. With this method, fibrin stains red, erythrocytes yellow, and connective tissue blue.

### Role of SVMPs in the thrombotic effect

Once the experimental model of thrombotic effect was established, the role of SVMPs in the pathogenesis of this effect was assessed by using the metalloproteinase inhibitor Batimastat (British Biotech, Oxford, UK). Solutions of venom from adult or juvenile specimens were prepared in PBS and incubated for 30 min at room temperature with Batimastat (final concentration 250 µM). Solutions of venoms incubated with the vehicle alone were also prepared. Aliquots of 100 µl of the mixtures, containing 70 µg venom, were injected i.p. into groups of four mice (22-23 g). Four hr after injection, mice were sacrificed and tissue samples from the lungs were obtained and processed for histological observation, as described above.

### Statistical analyses

Results were expressed as mean ± SEM. The significance of the differences between the mean values of experimental groups was assessed by Mann-Whitney U test when two groups were compared. In experiments involving more than two groups the significance of the differences was assessed by one-way ANOVA for normally distributed data and by Kruskal-Wallis test for non-normally distributed data. Tukey-Kramer or Dunn’s post-hoc tests, respectively, were used to analyze differences between pairs of mean values. P values < 0.05 were considered significant.

## Results

### Electrophoretic and chromatographic analyses

SDS-PAGE separation of venom revealed a highly similar electrophoretic pattern (Fig 1), with bands in the range of 150 kDa to 10 kDa. The most abundant bands have estimated molecular masses of 50, 30, 25, 22 and 15 kDa. A qualitatively similar RP-HPLC profile was also observed in these venoms (Fig 2), with few differences in peaks eluting at 52 min (juvenile venom) and 58, 65 and 83 min (adult venom).

**Figure 1:**
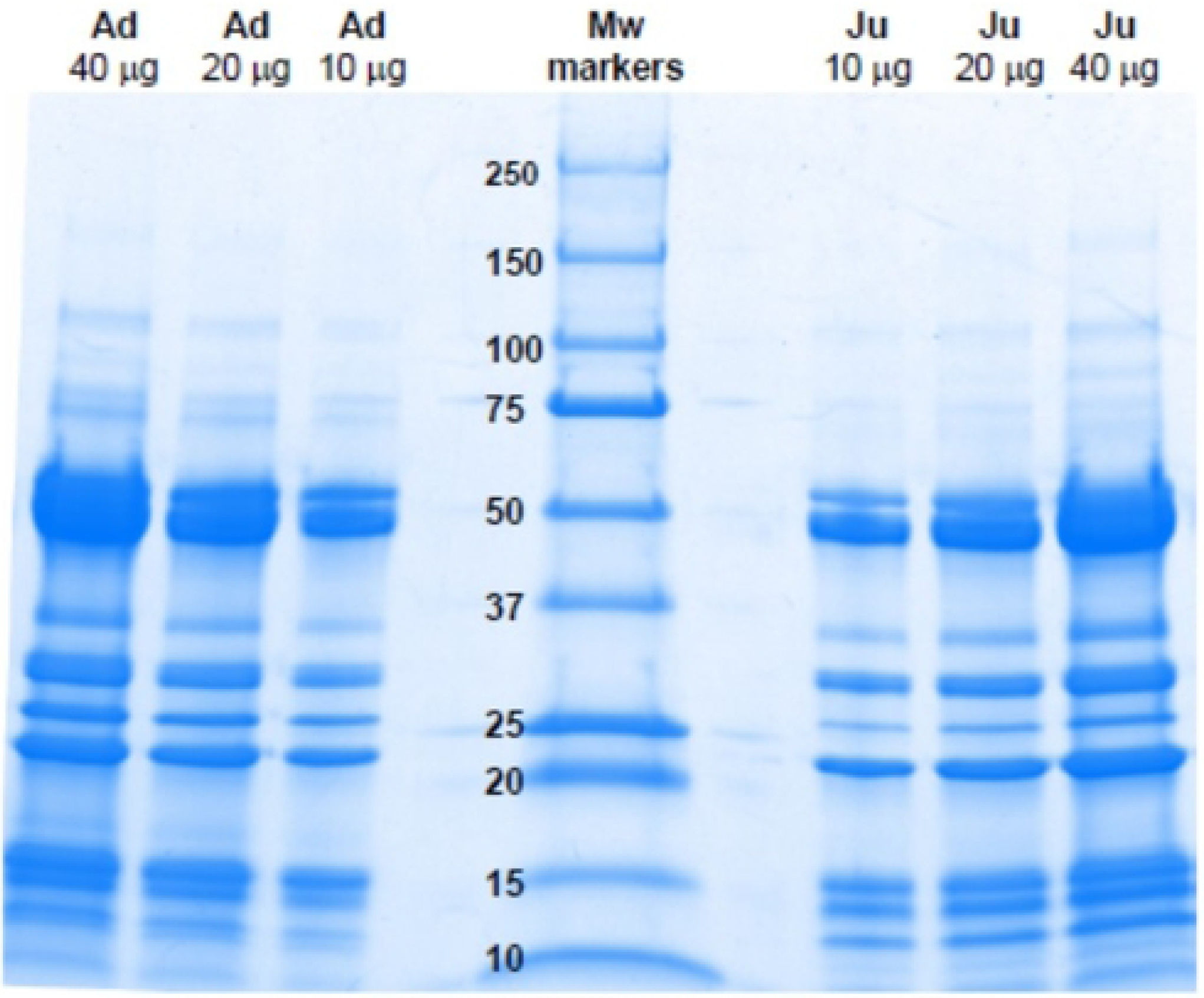
SDS-PAGE separation of venoms of adult (Ad) and juvenile (Ju) specimens of *B. lanceolatus*. Various amounts of venom (10, 20 and 40 µg) were separated under reducing conditions on 4-20% pre-cast gels. Molecular weight (Mw) markers were also run. Proteins were stained with Coomassie Blue R-250.

**Figure 2:**
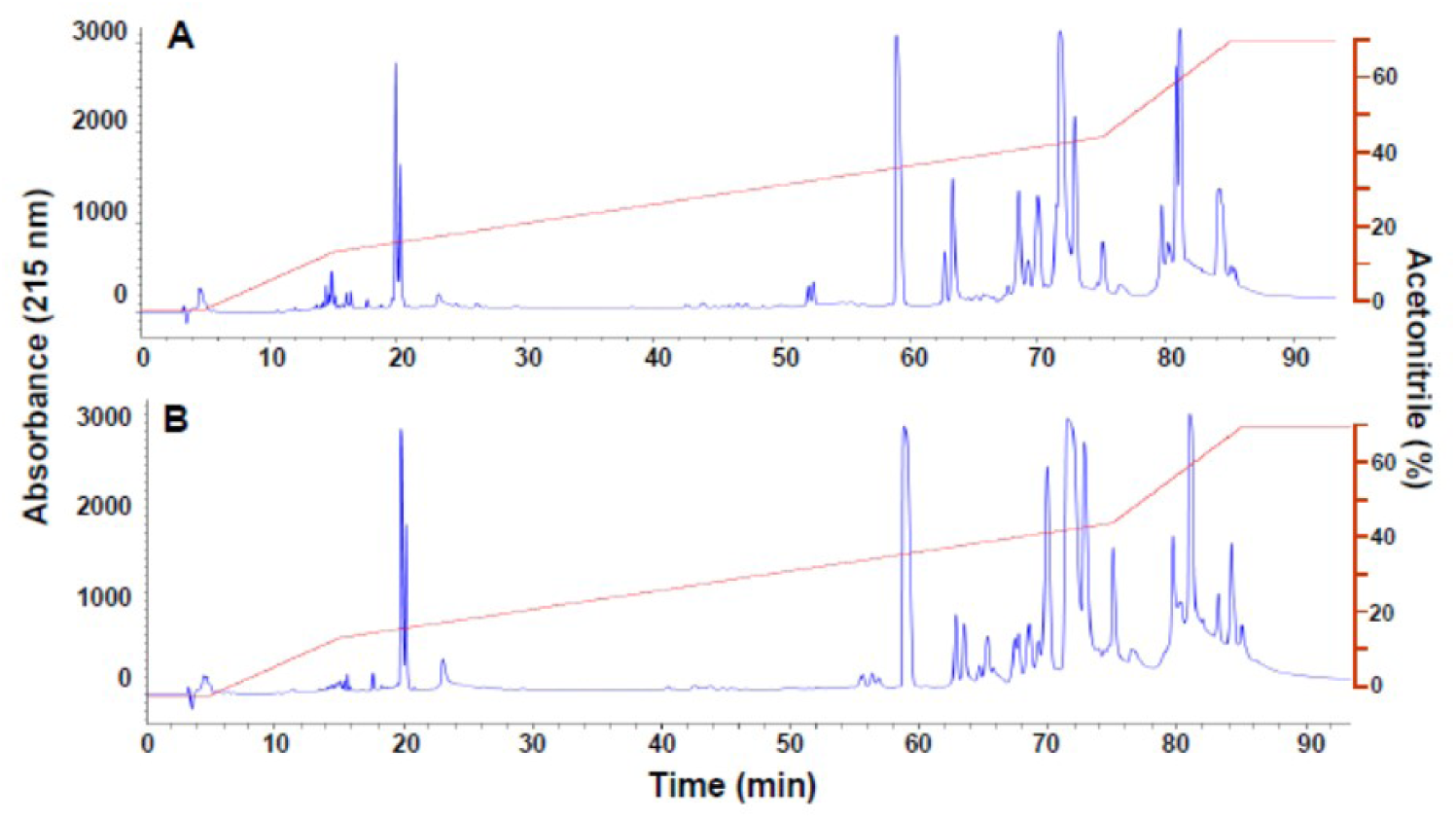
RP-HPLC separation of venoms of juvenile (A) and adult (B) specimens of *B. lanceolatus*. Samples of 2 mg were dissolved in water containing 0.1% trifluoroacetic acid (solution A). After centrifugation, the supernatant was applied to a reverse-phase column equilibrated with solution A and separation was monitored by recording the absorbance at 215 nm. The following gradient of acetonitrile, containing 0.1% trifluoroacetic acid, was used for elution: 0% for 5 min, 0-15% for 10 min, 15-45% for 60 min, 45-70% for 10 min, and 70% for 9 min (red line).

### Proteomic analysis

Table 1 depicts the protein families identified in the venoms of *Bothrops lanceolatus* (adults and juveniles) by bottom-up shotgun proteomics. Families present in both venoms include metalloproteinases (SVMPs), serine proteinases (SVSPs), phospholipases A_2_ (PLA_2_), C-type lectin-like proteins, L-amino acid oxidases, nerve growth factor, phospholipase B, and glutamyl cyclase, with highest number of variants in the first four families (Table 1). On the other hand, nucleotidase, vascular endothelial growth factor, hyaluronidase and PLA_2_ inhibitor were detected only in the adult venom, whereas protein disulfide isomerase was detected only in juvenile venom (Table 1). Supplementary Table S1 provides the details of the protein matches and supporting peptides.

**Table 1:**
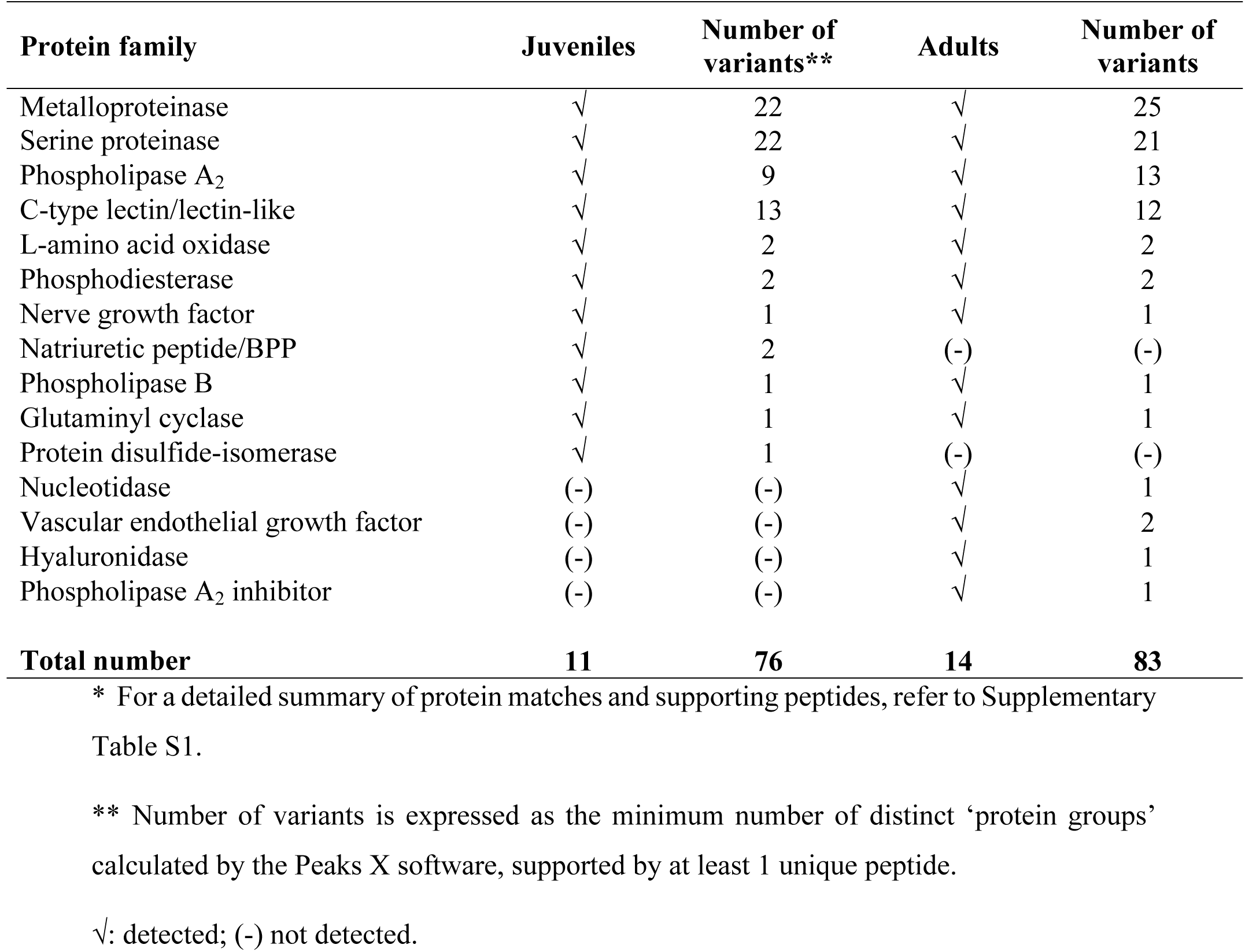
Protein families identified in the venoms of *Bothrops lanceolatus* (adults and juveniles) by bottom-up shotgun proteomics*.

### Procoagulant activity on plasma and fibrinogen *in vitro*

When added to citrated plasma in the presence of calcium, the venom of juvenile specimens of *B. lanceolatus* did not induce the formation of a firm clot up to a dose of 50 µg for a maximum observation period of 10 min, although an increase in turbidity was observed. When testing the venom of adult specimens, a weak clot formed after approximately 10 min of incubation when using a dose of 50 µg venom, whereas no clot formation, and only an increase in turbidity, occurred when using doses of venom of 25 µg, 12.5 µg, and 6.25 µg.

When venoms were added to a bovine fibrinogen solution, the venom of adult specimens induced the formation of a fibrin clot in a dose-dependent way. The time required to form a visible clot was 10 min, 9 min, 4.5 min and 3 min for venom amounts of 6.25 µg, 12.5 µg, 25 µg and 50 µg, respectively. In contrast, the venom of juvenile specimens did not induce the formation of a firm fibrin clot when the same doses were tested, although an increase in turbidity was observed at the dose of 50 µg. When the more sensitive turbidimetric method was used, both venoms induced an increase in absorbance at 340 nm, reflecting the formation of fibrin after addition of venoms to plasma and fibrinogen, thus revealing a thrombin-like (pseudo-procoagulant) activity (Fig 3). Venom of adult specimens showed a stronger thrombin-like activity than venom of juvenile specimens.

**Figure 3:**
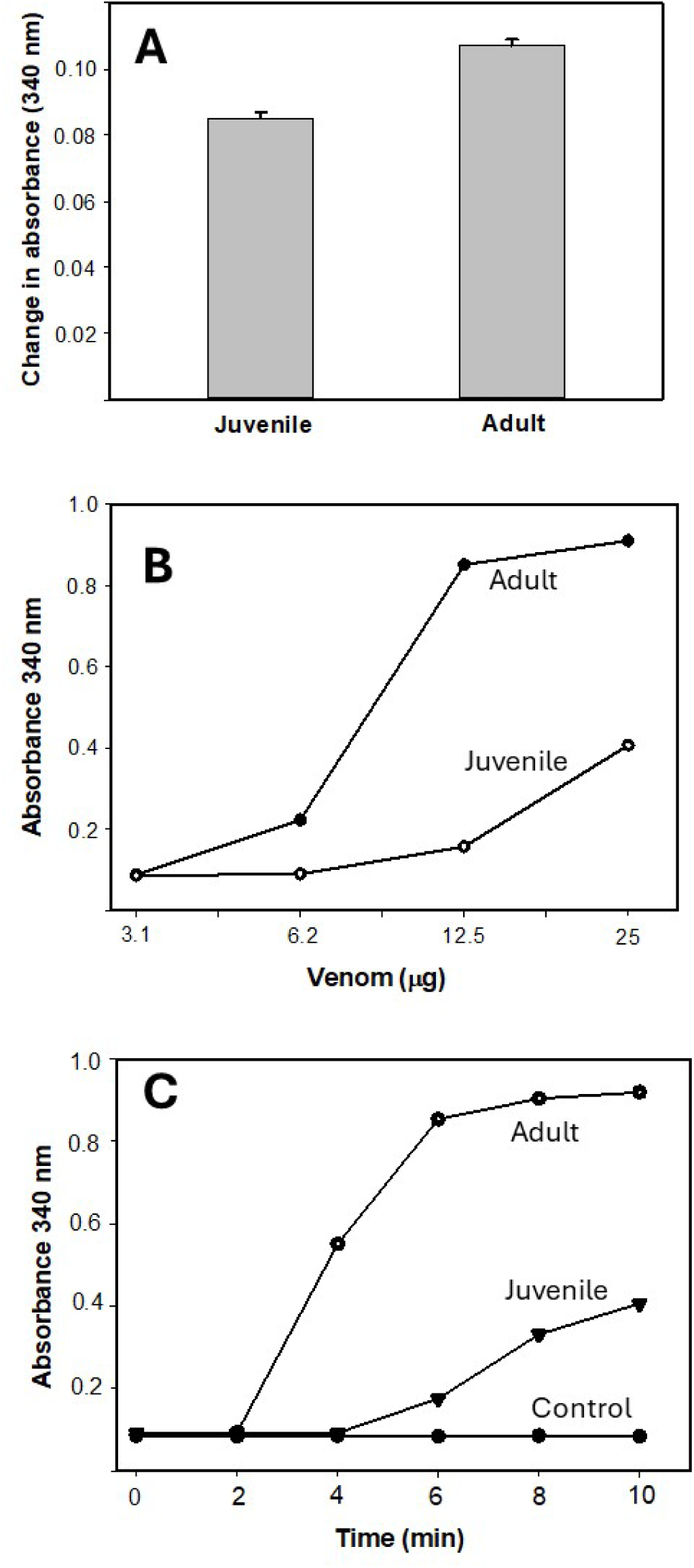
*In vitro* procoagulant activity of venoms of juvenile and adult specimens of *B. lanceolatus*. A: Procoagulant effect on plasma. Fifty µg venom, dissolved in 100 µL TBS, were added to wells in a microplate. Then, 4 µL of 0.4 M CaCl_2_ were added to 100 µL of mouse citrated plasma previously incubated at 37°C, and the mixture added to wells in the plate containing the venom solution. Controls included plasma/CaCl_2_ incubated with TBS with no venom. After shaking for 5 sec, the absorbances at 340 nm were recorded during 10 min as an index of the increase in turbidity of the samples. Results are presented as mean ± SEM (n = 3); p < 0.05 by Mann-Whitney U test when comparing the venoms. B and C: Thrombin-like (pseudo-procoagulant) activity effect of venoms on bovine fibrinogen. In B, solutions containing various amounts of venom, dissolved in 100 µL TBS, were added to 100 µL of a fibrinogen solution (6 mg/mL) previously incubated at 37°C and the change in absorbance at 340 nm were recorded at 10 min. In C, solutions containing 25 µg venom, dissolved in 100 µL TBS, were added to 100 µL of fibrinogen (6 mg/mL). The changes in absorbance at 340 nm were recorded at various time intervals. Assays were run in triplicates.

### Effect of clotting parameters *in vivo*

Experiments were done in order to assess the effect of venoms of juvenile and adult specimens on classical clotting tests and rotational thromboelastometry parameters. For this, two experimental settings were used: i.v. injection of 20 µg venom followed by bleeding and testing one hr after injection, and i.p. injection of 70 µg venom followed by bleeding and testing at 4 hr, the time when thrombosis was observed in pulmonary blood vessels. When the i.v. route was used, no major changes were observed in PT and aPTT in envenomed mice receiving either venom, as compared to control mice receiving PBS. However, a partial and significant drop in fibrinogen concentration occurred at one hr after i.v. injection of both venoms (Fig 4).

**Figure 4.**
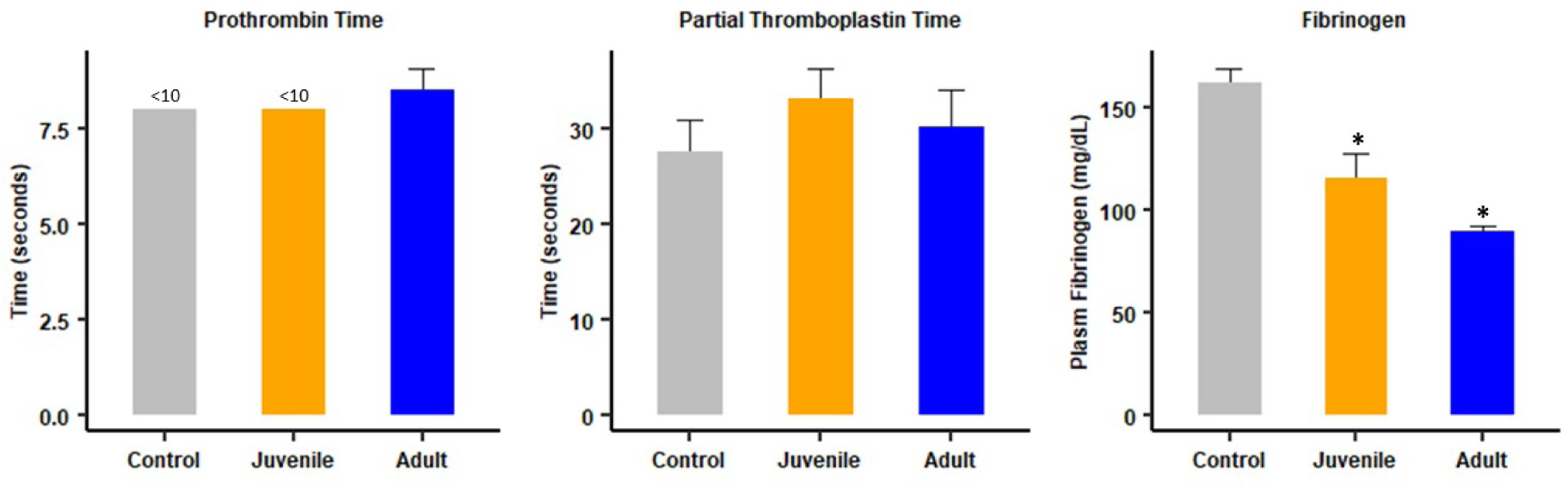
Effect of i.v. injection of juvenile and adult *B. lanceolatus* venoms on classical clotting tests. 20 µg venom from juvenile or adult specimens, dissolved in 100 µL PBS, were injected i.v. in mice. Controls received 100 µl PBS. One hour after injection mice were bled under isoflurane anesthesia and blood was collected, added to citrate anticoagulant, and centrifuged for plasma collection to determine prothrombin time (PT), activated partial thromboplastin time (aPTT) and fibrinogen concentration. Results are presented as mean ± SEM (n = 4). *p < 0.05 when compared to control.

Regarding rotational thromboelastometry parameters in samples obtained 1 hr after i.v. injection, Extem, Intem and Fibtem CT and CFT were prolonged in mice injected with juvenile venom, whereas only Fibtem CT and CFT were prolonged in the case of adult venom. When the clot strength was assessed by the determination of the A20 parameter, it was altered in Extem, Intem and Fibtem in the case of juvenile venom, whereas only the Extem A20 was altered in the case of adult venom (Figs 5 and 6). Thus, overall, the venom of juvenile specimens induced more pronounced alterations in these parameters than the venom of adult specimens when injected by the i.v. route.

**Figure 5.**
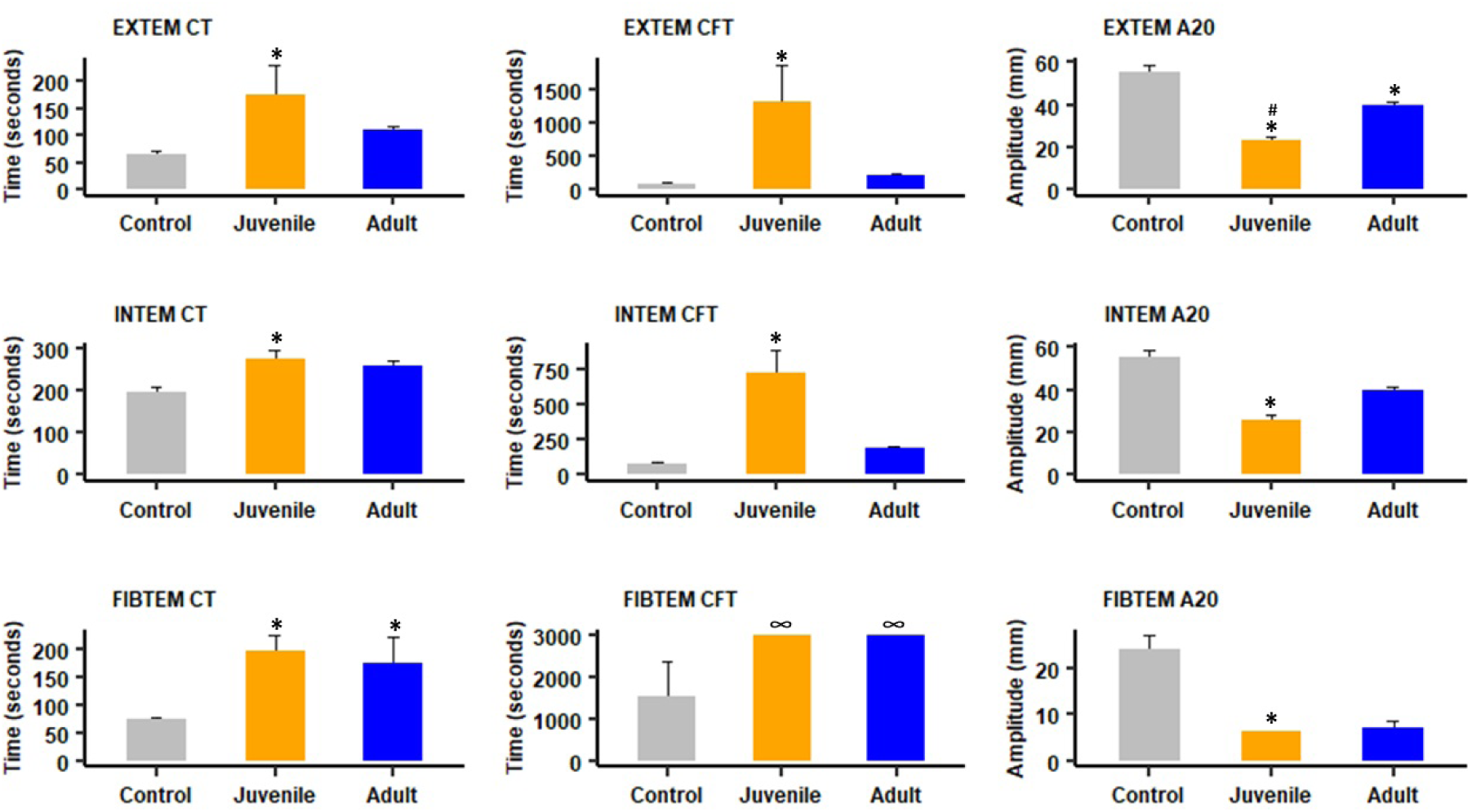
Effect of i.v. injection of juvenile and adult *B. lanceolatus* venoms on rotational thromboelastometry parameters. Twenty µg venom, dissolved in 100 µL PBS, was injected i.v. in mice. Controls received 100 µl of PBS. On hour after injection mice were bled under isoflurane anesthesia and blood was collected and added to citrate anticoagulant for determination of Extem, Intem and Fibtem parameters (see methods for details). Results are presented as mean ± SEM (n = 4). *p < 0.05 when compared to control; # p < 0.05 when comparing juvenile and adult venoms. In the case of Fibtem CFT in envenomed mice, no clot was formed.

**Figure 6.**
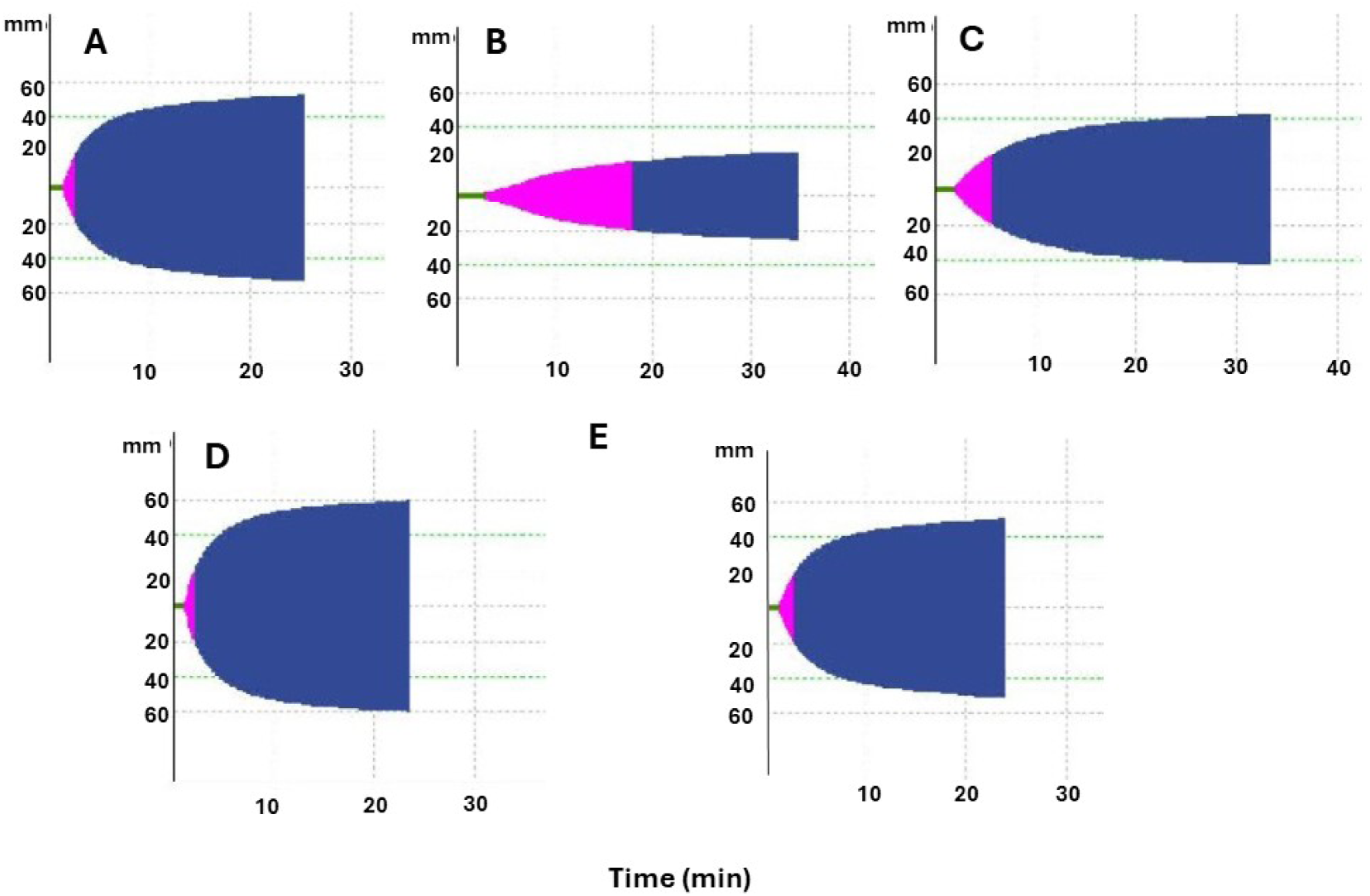
Representative rotational thromboelastometry tracings from mice injected with PBS or venoms of juvenile and adult specimens of *B. lanceolatus*. Mice were bled by cardiac puncture under isoflurane anesthesia, blood was added to sodium citrate solution, and evaluated by rotational thromboelastometry (see methods for details). (A) Extem tracing from a mouse receiving PBS by the i.v. route and bled 1 hr after injection. (B) and (C) Extem tracings from mice receiving 20 µg of juvenile (B) or adult (C) *B. lanceolatus* venoms by the i.v. route and bled 1 hr after injection. (D) and (E) Extem tracings from mice receiving 70 µg of juvenile (D) or adult (E) venoms by the i.p. route and bled 4 hr after injection.

In order to assess the status of clotting parameters in circumstances when pulmonary thrombosis occurred, the same assays were carried out in samples collected 4 hr after i.p. injection of 70 µg venom, which induces thrombosis. No alterations in PT, aPTT and fibrinogen concentration were observed in mice receiving adult venom, whereas a partial, but significant, change occurred in aPTT and fibrinogen in mice receiving juvenile venom (Fig 7). On the other hand, both venoms induced a profound thrombocytopenia (Fig 7).

**Figure 7.**
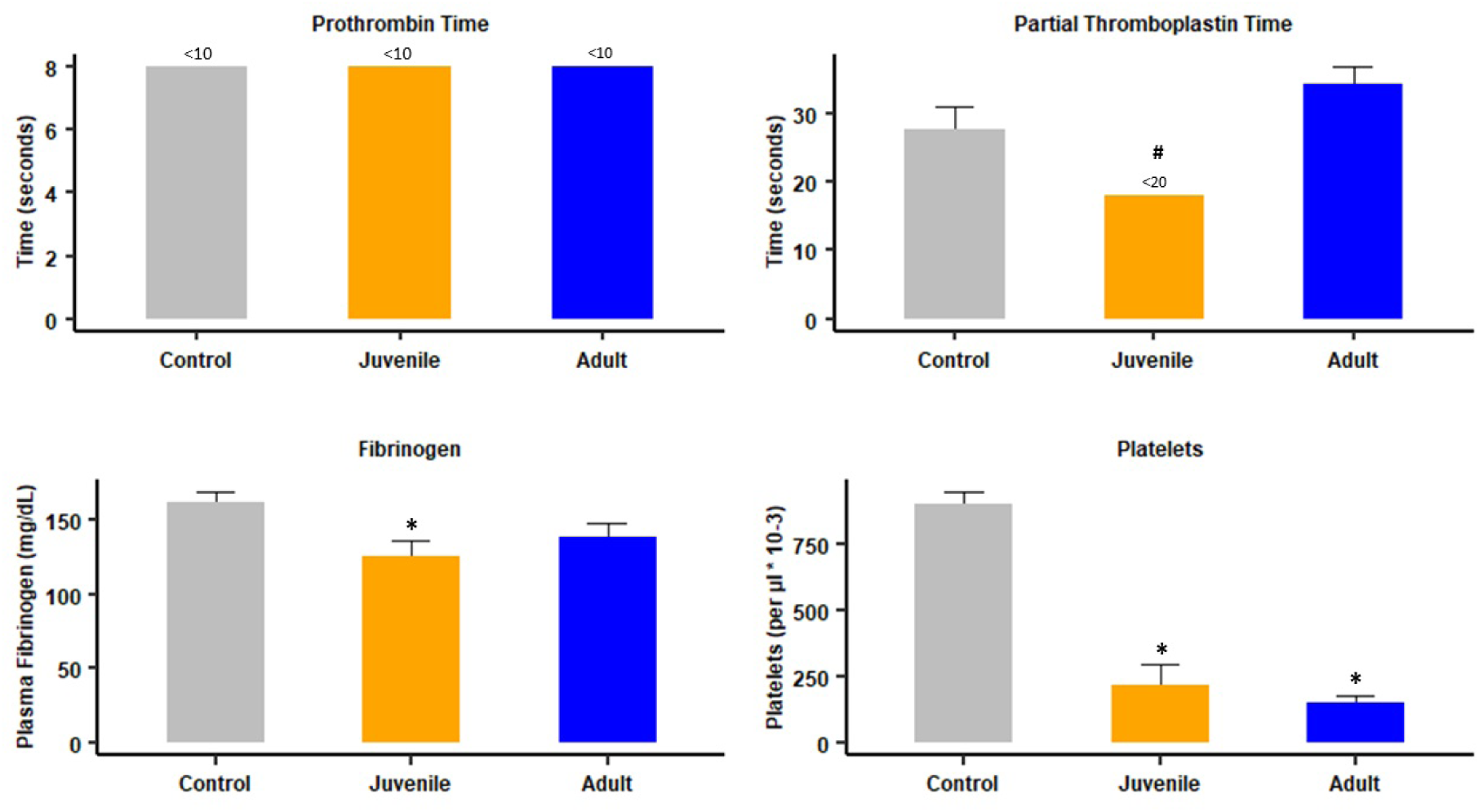
Effects of i.p. injection of juvenile and adult *B. lanceolatus* venoms on classical clotting tests and platelet counts. Mice received 70 µg venom, dissolved in 100 µL PBS, by the i.p. route. Controls received 100 µl of PBS. Four hours after injection mice were bled under isoflurane anesthesia and blood was collected and added to citrate anticoagulant for determination of prothrombin time (PT), activated partial thromboplastin time (aPTT), fibrinogen concentration, and platelet counts (see methods for details). Results are presented as mean ± SEM (n = 4 in the case of clotting tests and fibrinogen concentration and n = 4-11 in the case of platelet counts). *p < 0.05 when compared to control; # p < 0.05 when comparing juvenile and adult venom.

Analysis of rotational thromboelastometry parameters in blood samples collected 4 hr after i.p. injection of venoms showed no significant alterations in samples from envenomed mice, as compared to those from control mice (Fig 8). The raw data related to coagulation, rotational thromboelastometry and platelet counts are available in Supplementary Table S2.

**Figure 8.**
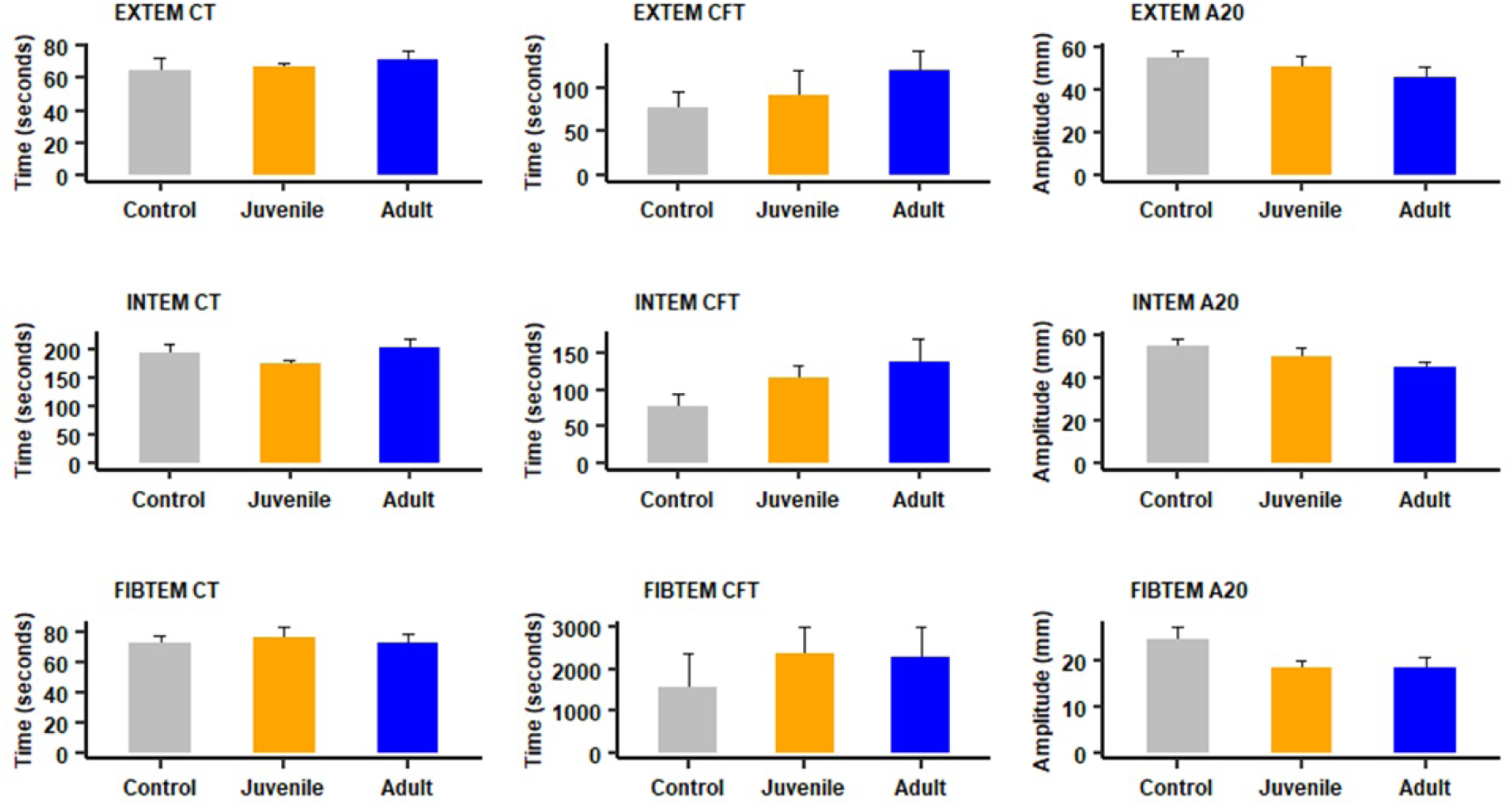
Effect of i.p. injection of juvenile and adult *B. lanceolatus* venoms on rotational thromboelastometry parameters. Mice received 70 µg of venom, dissolved in PBS, by the i.p. route. Controls were injected with 100 µl of PBS. Four hours after injection mice were bled under isoflurane anesthesia and blood was added to citrate anticoagulant for determination of Extem, Intem and Fibtem parameters (see methods for details)]. Results are presented as mean ± SEM (n = 4).

### Histological assessment of thrombosis

Tissue samples from control mice receiving injections of PBS by various routes showed normal histological features in heart, brain, and lungs (Fig 9). In the cases of animals receiving venoms, no thrombi were observed in brain, heart, and lung blood vessels after adult and juvenile *B. lanceolatus* venom injections by the i.v., s.c. and i.m. routes at 4 hr and 24 hr. In contrast, when 70 µg venom were administered by the i.p. route, the venom of juvenile *B. lanceolatus* specimens induced numerous thrombi in the lungs at both time intervals (Fig 9), but not in heart or brain. Few mice receiving 70 µg juvenile venom by the i.p. route died before 4 hr; in these cases, additional mice were injected to complete the sample of four mice per time interval. In samples from mice receiving juvenile venom by the i.p. route thrombi were observed in vessels of variable caliber, being more abundant in veins, although they were also present in arteries and arterioles. On the other hand, venom of adult specimens induced the formation of thrombi only in few blood vessels and in few of the tissue samples analyzed, and no thrombi were observed in brain and heart vasculature. In some pulmonary vessels, mostly veins, of mice injected with adult venom a hyaline pale material was observed having a lower density as compared to the overt thrombi observed with juvenile venom (Fig 9). No mice injected with adult venom by the i.p. route died.

**Figure 9:**
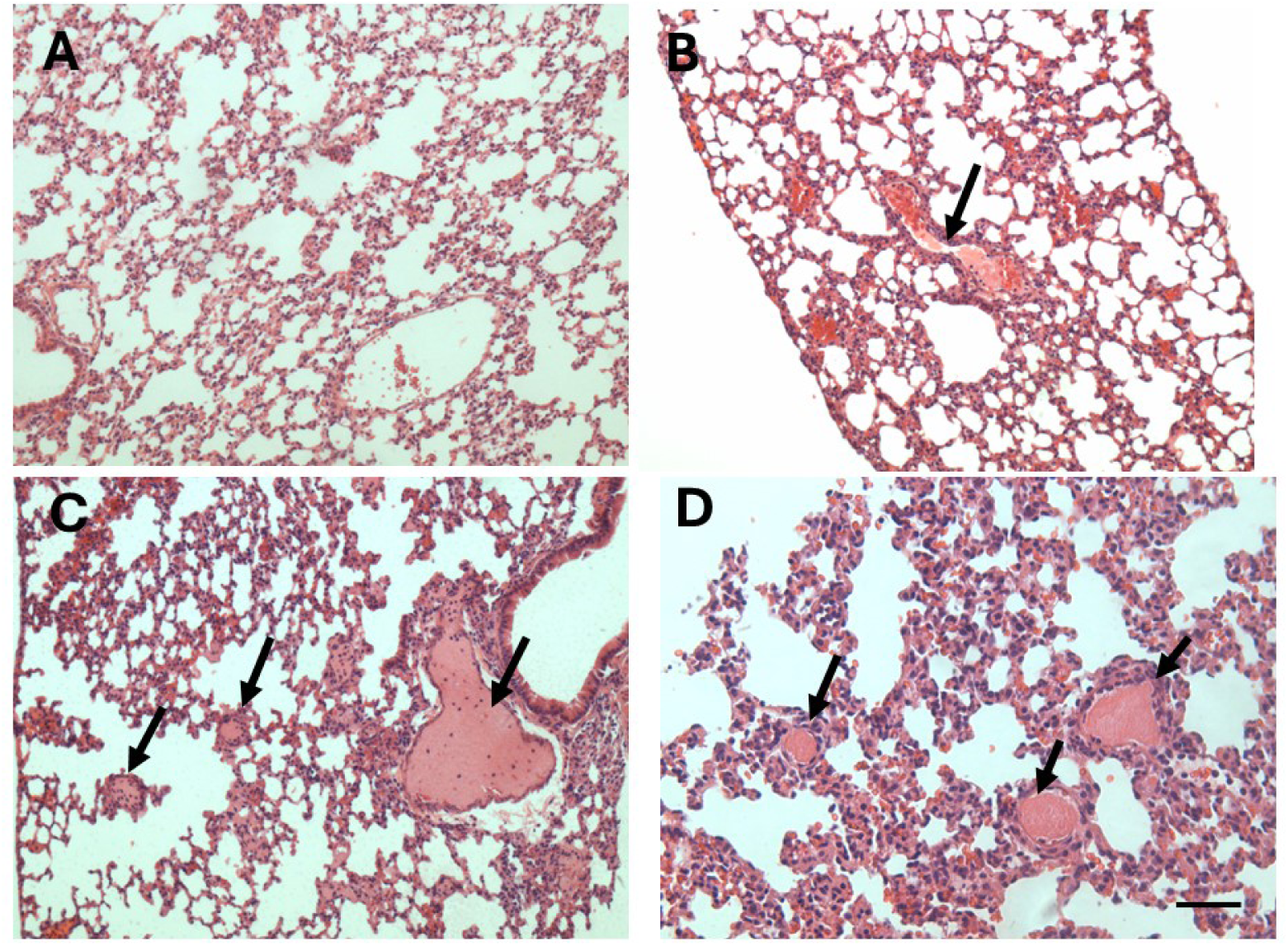
Light micrographs of sections of pulmonary tissue of mice which received an i.p. injection of venom from juvenile or adult *B. lanceolatus*. Sections correspond to samples of mice receiving 100 µL PBS (A) or 70 µg of *B. lanceolatus* venom from either adult (B) or juvenile (C, D) specimens, dissolved in 100 µL PBS. Mice were sacrificed 4 hr after injections, and samples from pulmonary tissue were collected, added to formalin fixative and processed for embedding in paraffin. Sections from control mice receiving PBS show a normal histological pattern. In sections of mice receiving adult venom, thrombi are largely absent and only a hyaline material is present in some vessels (arrow). In contrast, samples from mice injected with juvenile venom present abundant thrombi in veins, arteries, and smaller blood vessels (arrows). Hematoxylin-eosin staining. Bar represents 100 µm.

Overall, blood vessels with thrombi were more abundant in mice injected with juvenile venom than in those receiving adult venom. In order to provide a semiquantitative assessment of the frequency of thrombi formation, a number of histological sections obtained from mice injected with venoms were examined. Among 26 sections from mice receiving juvenile venom, 20 showed thrombi, 3 presented the hyaline pattern of staining described above, and 3 did not show thrombi. In contrast, in the case of 16 sections from lung tissue of mice injected with adult venom, 1 had thrombi, 6 presented the hyaline material inside vessels, and 9 did not show thrombi or hyaline material. It was of interest to assess whether thrombi were also present in the microvasculature of mice receiving juvenile venom, by using a specific staining of fibrin (Martius-Scarlet-Blue). No red staining, characteristic of fibrin, was observed in the microvasculature of alveolar septa in samples from mice injected with PBS, whereas abundant red-stained microthrombi occurred in sections from mice that had received an i.p. injection of venom from juvenile specimens 4 hr after injection (Fig 10).

**Figure 10.**
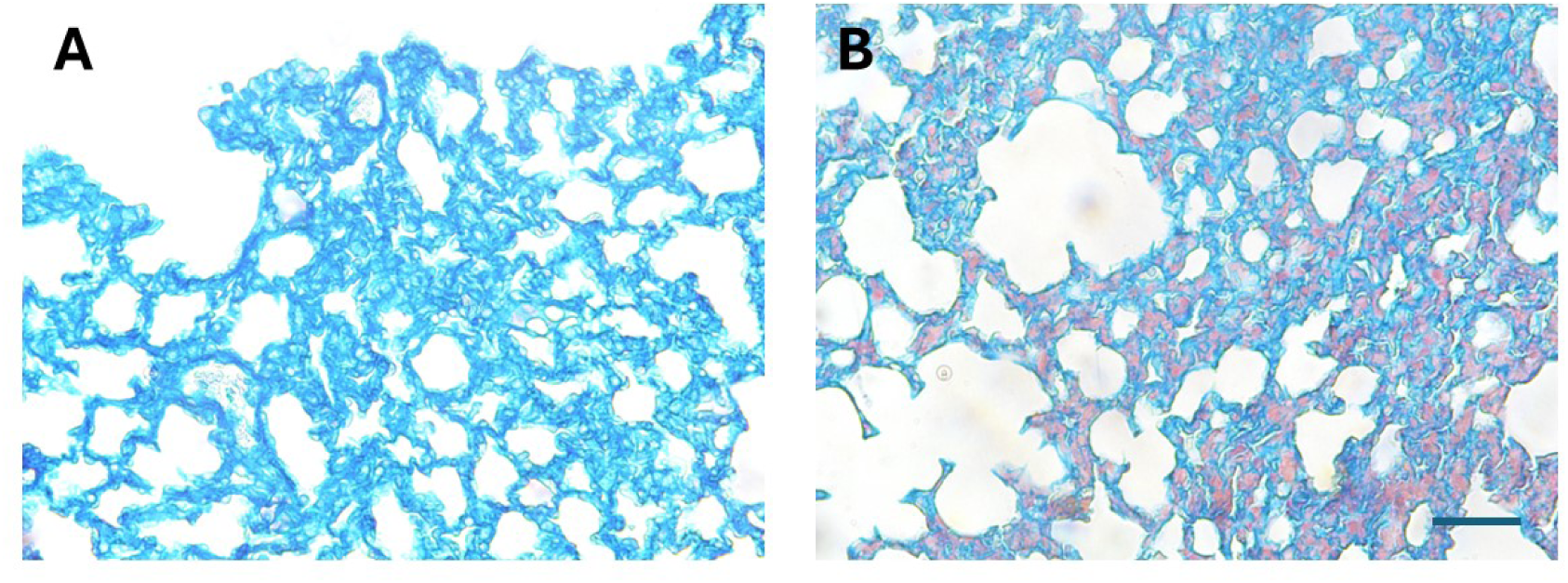
Thrombi in the pulmonary microvasculature of mice injected with juvenile *B. lanceolatus* venom. Light micrographs of sections of pulmonary tissue of mice which received an i.p. injection of 100 µL PBS (A) or 70 µg of *B. lanceolatus* venom from juvenile specimens, dissolved in 100 µL PBS (B). Mice were sacrificed 4 hr after injections, and samples from pulmonary tissue were collected, added to formalin fixative and processed for embedding in paraffin. Sections were stained with Martius-Scarlet-Blue, which stains fibrin in red color. No fibrin is observed in samples of mice receiving PBS, whereas abundant red fibrin deposits are observed in the microvasculature of the section from mice treated with venom. Bar represents 100 µm.

In order to assess the role of SVMPs in the formation of pulmonary thrombi, venoms were incubated with the metalloproteinase inhibitor Batimastat before i.p. injection. Such treatment did not inhibit the formation of thrombi in samples injected with juvenile venom, and also did not prevent the formation of thrombi or the hyaline intravascular material in the case of adult venoms. In contrast, Batimastat was effective in the inhibition of hemorrhagic effect in the lungs (Fig 11). Thus, SVMPs do not seem to be the causative agent of pulmonary thrombosis in this model but are responsible for the hemorrhagic activity. The raw data related to histological assessment of thrombosis are available in Supplementary Table S3.

**Figure 11.**
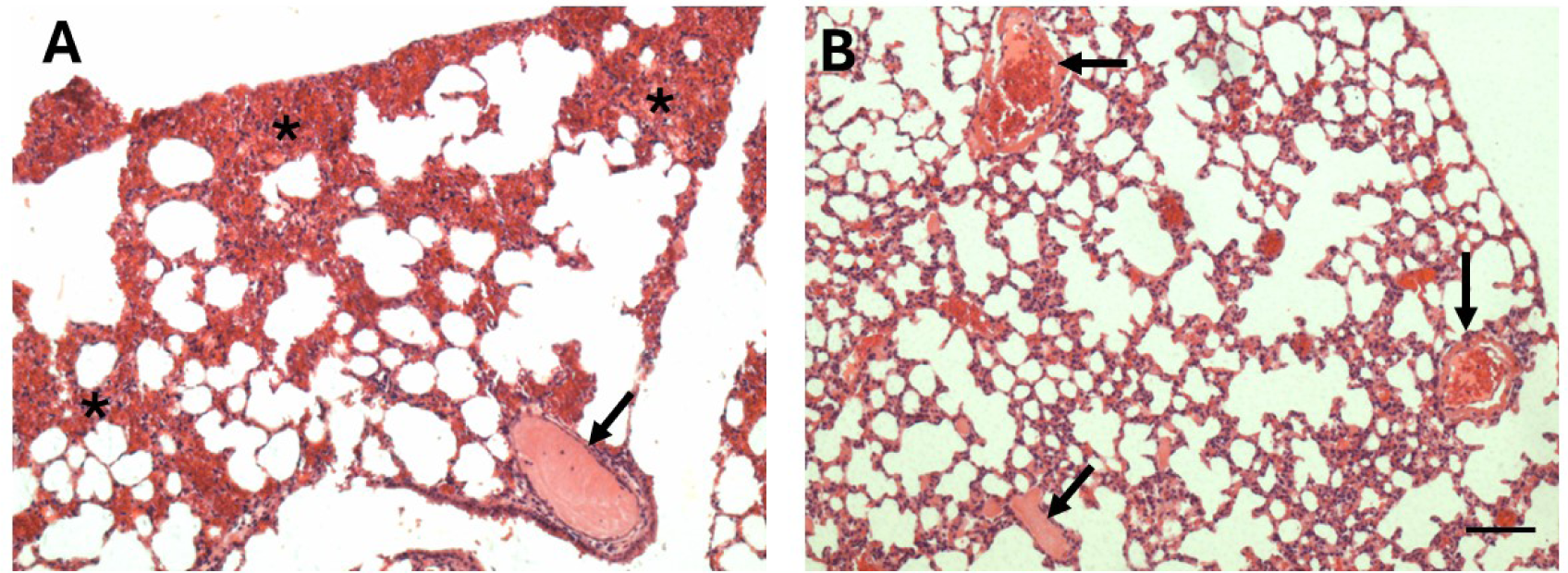
Inhibitory action of Batimastat on the effects induced by juvenile *B. lanceolatus* venom in the lungs. Light micrographs of sections of pulmonary tissue of mice which received an i.p. injection of 70 µg of *B. lanceolatus* venom from juvenile specimens dissolved in 100 µL PBS (A) or the same dose of venom which was previously incubated with Batimastat (250 µM final concentration) (B). Mice were sacrificed 4 hr after injections, and samples from pulmonary tissue were collected, added to formalin fixative and processed for embedding in paraffin. The section from a mouse receiving venom show abundant hemorrhage (*) and a thrombus in a blood vessel (arrow). In contrast, no hemorrhage is observed in tissue from a mouse receiving venom incubated with Batimastat, whereas thrombi are present (arrows). Hematoxylin-eosin staining. Bar represents 100 µm.

## Discussion

The mouse experimental model described in this work reproduces three of the characteristic clinical findings in patients envenomed by *B. lanceolatus*, i.e., thrombosis, lack of consumption coagulopathy, and thrombocytopenia. Clinical observations indicate that patients developing thrombosis were generally bitten by specimens of small size [4, 22]. This prompted us to obtain venoms of juvenile snakes and compare their composition and effects with those of adult specimens. Results show that, when administered by the i.p. route, venom from juvenile specimens induced pulmonary thrombosis in mice, whereas the venom of adult specimens induced this effect to a much lower extent. Interestingly, no such effect was observed when venom was injected i.v., s.c. or i.m. When administered by the i.p. route, macromolecules reach the systemic circulation mainly via lymphatic vessels and reach higher concentrations in blood than when using the s.c. and i.m. routes [31]. It is suggested that the i.p. route is a model of systemic absorption of the venom that, in terms of toxicokinetics, may resemble what occurs in clinical cases. Thus, our model could be used to study of the mechanism of thrombosis by *B. lanceolatus*.

The majority of thrombi observed in pulmonary vasculature in mice injected with juvenile venoms were observed in large and medium size veins, although thrombi were also present in arteries. In addition, a staining for fibrin showed positive staining in the microvasculature of the alveolar septa, hence reproducing the phenomenon of diffuse thrombotic microangiopathy described in an autopsy of a patient envenomed by *B. lanceolatus* [12]. In our study we did not observe thrombi in cerebral or myocardial blood vessels, which are common findings in human patients [2–4]. Our observations can be explained by the higher propensity of the pulmonary vasculature to develop inflammatory and thrombotic complications in a variety of diseases [32, 33] and by its unique hemodynamic and immunologic features. In the case of tissue samples from mice injected with adult venoms the occurrence of evident thrombi was infrequent. Instead, a pale hyaline material was observed in some vessels. Clearly, the thrombotic action induced by adult venoms was weaker than by juvenile venoms.

The proteomic analysis of juvenile and adult venoms did not reveal overt differences in composition. Thus, *B. lanceolatus* presents a venom proteomic profile that fits within the ‘paedomorphic’ pattern described for some populations of *B. atrox*, in which there are no major changes in venom composition as snakes age [34]. *Bothrops* sp venoms present a dichotomic ontogenetic pattern, with some venoms having a ‘paedomorphic’ profile while others show drastic changes as snakes age, corresponding to an ‘ontogenetic’ profile [34,35]. Thus, the basis for the difference in thrombogenic potential between juvenile and adult venoms is not evident from the overall electrophoretic, chromatographic, and proteomic comparison. It is likely that variations in the action of toxins within the most abundant protein families may account for the functional difference observed, an issue that awaits the identification of the thrombogenic factor(s) in venoms of juvenile specimens. The low amount of venom available from juvenile specimens did not allow us to undertake the purification of venom components.

The use of the SVMP inhibitor Batimastat allowed us to assess whether SVMPs are involved in the thrombotic effect. It has been hypothesized that *B. lanceolatus* venom may activate the vascular endothelium, rendering it thrombogenic [12, 36], an effect that might be related to the action of SVMPs, which are abundant in this venom [14, 19, 37, 38]. It is known that SVMPs induce a variety of effects on endothelial cells [39–41]. However, inhibition of SVMPs did not prevent the formation of thrombi, although abrogated pulmonary hemorrhage, thus evidencing that thrombosis occurs in conditions where SVMPs are inhibited and implying that other as yet unidentified components are responsible for this effect. The proinflammatory effect of *B. lanceolatus* venom, reflected by its ability to activate the complement system and generate a variety of mediators, has been proposed as a possible mechanism of the thrombotic effect [16–18]. Alternative mechanisms of thrombosis might be related to the action of venom on von Willebrand factor, promoting its binding to type VI collagen in the subendothelial surface [15] or to platelet activation, perhaps associated with the thrombocytopenia observed in clinical cases in in our experimental conditions. Our observations concur with clinical laboratory findings in that this venom induces thrombocytopenia, which might be a consequence of the action of a C-type lectin-like component, similar to the one characterized from the closely related venom of *B. caribbaeus* [42]. It is likely that the pathogenesis of thrombosis involves an interplay between inflammatory events and platelet alterations in a scenario of thromboinflammation [43, 44].

There have been conflicting findings in the literature concerning the effect of *B. lanceolatus* venom on hemostasis *in vitro* and *in vivo*. Regarding the *in vitro* procoagulant effect on citrated plasma, both negative and positive results have been described [14, 19, 21,23, 24], whereas the venom did not induce defibrinogenation in mice [21]. It has been proposed that the negative *in vitro* results described by Bogarín et al [21] and Gutiérrez et al. [14] are due to the fact that calcium, a cofactor required for coagulation, was not added to the plasma in these experiments. When calcium is added, adult *B lanceolatus* venom induces plasma clotting *in vitro* [19, 23, 24]. Our observations and those of others demonstrate that venoms of both adult and juvenile specimens exert thrombin-like (pseudo-procoagulant) activity. This activity has been previously described for adult *B. lanceolatus* venom [20, 24]. This effect explains the weak procoagulant activity described for this venom on citrated plasma. However, *in vitro* experiments do not necessarily reproduce what occurs *in vivo* and hence the importance of assessing hemostatic alterations *in vivo*.

We explored the *in vivo* effect of adult and juvenile venoms on classical clotting tests and rotational thromboelastometry parameters. No alterations were observed in PT and aPTT, and there was only a significant drop in the concentration of fibrinogen after i.v. injection. In contrast, when the venom of *B. asper*, which induces a typical consumption coagulopathy, was tested in a similar mouse model, there were drastic alterations in these parameters [45]. This agrees with clinical observations of *B. lanceolatus* envenomings since, in most cases, a consumption coagulopathy is not observed [3, 5, 11].

When *in vivo* effects were evaluated by rotational thromboelastometry, which provides a more detailed assessment of the hemostatic status, the venom of juvenile specimens affected various Extem, Intem and Fibtem parameters 1 hr after i.v. injection, whereas only Fibtem parameters were altered in the case of adult venom. These findings might be due to the partial drop in fibrinogen concentration in mice injected with *B. lanceolatus* venoms as a consequence of the action of thrombin-like (pseudo-procoagulant) enzymes, since Fibtem basically depends on the status of fibrinogen. However, when venoms were administered i.p., in conditions where pulmonary thrombosis occurs, clotting tests were not altered, implying that thrombosis occurs in conditions where there is no defibrinogenation. The higher extent of rotational thromboelastometry alterations when using the i.v. route, as compared to the i.p. route, might be due to the fact that in the former the venom is in immediate contact with clotting factors in the bloodstream, thus increasing the likelihood of alterations. The lack of major alterations in rotational thromboelastometry parameters is in contrast with observations carried out with the venom of *B. asper*, which causes a consumption coagulopathy and drastically affects these parameters in a mouse model [45]. Overall, *B. lanceolatus* venoms do not induce *in vivo* consumption coagulopathy in this model, in agreement with clinical observations. The lack of overt consumption coagulopathy might be related to the thrombotic effect since clotting factors are present in the bloodstream by the time the thrombogenic mechanisms induced by the venom are operating in blood vessels. On the other hand, a drastic drop in platelet numbers was observed by the time thrombosis occurs, suggesting that platelet alterations might be related to the pathogenesis of thrombosis by mechanisms as yet unknown.

In conclusion, when injected by the i.p. route, the venom of juvenile specimens of *B. lanceolatus* induces abundant thrombi in the pulmonary vasculature, thus reproducing clinical findings describing thrombosis in patients bitten by snakes of small size in Martinique. The model also reproduces clinical observations describing lack of consumption coagulopathy and thrombocytopenia in many patients. This experimental model could be used to explore the mechanisms of thrombosis induced by the venom of *B. lanceolatus* and to identify the toxins responsible for this unique effect. Moreover, although rare, peripheral arterial thrombosis and pulmonary thromboembolism have been described in envenomings by other viperid snake species [46, 47], thus opening the possibility of using this experimental model to explore the development of thrombosis with other snake venoms.

## Acknowledgments

The authors thank Marilla Lamela Méndez (Capris S.A., Costa Rica) for providing the reagents for the rotational thromboelastometry assays and Jennifer Stynoski for her collaboration in the statistical analyses. This study was supported by Vicerrectoría de Investigación, Universidad de Costa Rica (project 01-273-2024).

## Supporting information

**Supplementary Table S1**. Details of the protein matches and supporting peptides related to the proteomic analysis of venoms of *B. lanceolatus* from adult and juvenile specimens.

**Supplementary Table S2**. Raw data of the assays evaluating the alterations induced by the venoms of *B. lanceolatus* on classical clotting tests, rotational thromboelastometry, and platelet counts.

**Supplementary Table S3**. Raw data of the histological assessment of thrombosis in histological sections from the lungs of mice receiving intraperitoneal injection of 70 µg of the venoms of juvenile and adult specimens of *B. lanceolatus*.

